# Watering regime affects aboveground and belowground trait covariation in a hybrid *Silphium* population

**DOI:** 10.1101/2025.09.26.678844

**Authors:** Tyler Thrash, Meredith T. Hanlon, Matthew McNeese, Matthew J. Rubin, David L. Van Tassel, Allison J. Miller

**Author notes:** Manuscript received; revision accepted.

## Abstract

**Premise of the study:** Perennial, herbaceous members of the sunflower family (Asteraceae) can provide valuable agronomic products from aboveground biomass, as well as ecosystem services belowground. Aboveground and belowground trait covariation can be affected by environmental conditions such as the location of available water. Here, we investigated the effects of watering regime on trait covariation in *Silphium* (Asteraceae), an emerging perennial oilseed crop.

**Methods:** *Silphium integrifolium* Michx. and S. *perfoliatum* L. are native North American species adapted to well-drained and moist soils, respectively. Using a backcrossed F1 hybrid *S. integrifolium x S. perfoliatum* population, we conducted a greenhouse experiment where plants were either watered at the soil line (“top-watered”) or from below using watering trays (“bottom-watered”). Aboveground biomass, root distribution, and root morphology (e.g., specific root length) were measured destructively.

**Key results:** Compared to top-watered plants, bottom-watered plants were smaller aboveground and belowground, with a higher proportion of roots in the upper portions of the pots and higher specific root length. Negative relationships between aboveground biomass and specific root length were steeper for bottom-watered than top-watered plants.

**Conclusions:** The location of available water impacts aboveground biomass, the distribution and morphology of the root system, and relationships between aboveground and belowground plant structures. This work suggests that resource allocation depends in part on the location of available water, and has relevance for ongoing efforts aimed at developing perennial, herbaceous species that offer both economically viable products as well as ecosystem services.

## INTRODUCTION

Emerging perennial crops have the potential to provide seed, oil, and other economically important products aboveground, as well as ecosystem services such as carbon sequestration and reduced soil erosion belowground (Crews et al., 2018). Some perennial members of the sunflower family (Asteraceae) are currently undergoing domestication for oil seeds and biofuels (Van Tassel et al., 2017), including *Silphium integrifolium* Michx. (Price et al., 2022; Reinert et al., 2019; Vilela et al., 2018) and *S. perfoliatum* L. (Assefa et al., 2015; Gansberger et al., 2015). To establish the viability of these two species as crops and identify pathways for improvement, traits related to aboveground growth and productivity have been characterized for multiple years in different environments (Assefa et al., 2015; Greve et al., 2023; Kowalski, 2004; Price et al., 2022). Although *Silphium integrifolium* has been identified as being deep-rooted and drought-tolerant since at least 1935 (Weaver & Stoddart, 1935), the roots of *Silphium* species have only recently started to be investigated (e.g., Gonzalez-Paleo et al., 2023; Gonzalez-Paleo et al., 2024).

The distribution of roots, including root depth, is the product of genetics, the environment, and their interaction. Annual domesticated sunflower species are relatively deep-rooted compared to other annual crops such as maize (Comas et al., 2013; Debaeke et al., 2021). Environmental stressors that affect the distribution of sunflower roots include drought stress (Rauf & Sadaqat, 2007), osmotic stress (Masalia et al., 2018), and salt stress (Ma et al., 2021). Specifically, lack of available water tends to decrease root biomass and increase total root length (Masalia et al., 2018), although these effects may differ by genotype (Rauf & Sadaqat, 2007) and developmental stage (Ma et al., 2021). Members of the perennial *Silphium* genus, such as *S. integrifolium* and *S. perfoliatum,* have roots ranging from massive tap roots to finer root systems that are more typical of many dicotyledonous species (Clevinger & Panero, 2000). This variation is most likely due to evolution in response to the different environments where these species are commonly found. While the two species have at least somewhat partially overlapping ranges east of the Rocky Mountains in the continental United States (USDA, 2025), *S. integrifolium* tends to grow in drier environments where roots need to obtain water from deeper soil layers (Vilela et al., 2018), and *S. perfoliatum* tends to grow in wetter environments where water is often readily available near the soil surface (Schoo et al., 2017). *Silphium* populations derived from hybrids between these two species may exhibit a wide variety of root distributions and root morphological characteristics, as well as covariation among root traits and aboveground structures.

Understanding natural variation in root traits, as well as how root traits covary with aboveground portions of the plant, is particularly important in emerging perennial crops undergoing selection for aboveground traits because the goal is to enhance productivity aboveground (yield) while simultaneously maintaining or enhancing root structures that provide key ecosystem services belowground (Crews et al., 2018). Trait covariation typically refers to relationships among pairs of traits and reflects how plants allocate resources for growth, reproduction, and survival (Agrawal et al., 2010; Sgrò & Hoffmann, 2004). Positive covariation between a pair of traits often indicates that the two traits respond to the same environmental variable in similar ways (Agrawal et al., 2010). For example, aboveground biomass positively covaries with belowground biomass in switchgrass (*Panicum virgatum)*: larger plants are larger both aboveground and belowground (Chen et al., 2021). Alternatively, negative covariation between a pair of traits may indicate that the two traits result from directional selection on the same resource, resulting in tradeoffs (Agrawal, et al., 2010). Negative covariation between two traits without evidence for directional selection may result from the relationships between the two traits and confounded aspects of the environment (Knops et al., 2007). Studying relationships among aboveground and belowground traits is challenging because accurate measurements require destructive sampling. In the case of perennials, multi-year growth can add complexity, and experimental systems often lack genetic replication.

*Silphium integrifolium,* a perennial oilseed undergoing domestication, is emerging as a model system for studying aboveground and belowground resource allocation (Gonzalez-Paleo et al., 2023; Gonzalez-Paleo et al., 2024; Vilela et al., 2018). A recent study investigated nitrogen (N) allocation in *S. integrifolium*, and reported that N allocated to leaves can negatively covary with N allocated to the root crown at pre-anthesis (i.e., during mostly vegetative growth; Gonzalez-Paleo et al., 2023).

Furthermore, N allocated to seeds can negatively covary with N allocated to the root crown at maturity (i.e., during mostly reproductive growth; Gonzalez-Paleo et al., 2023). Similarly, selecting for early vigor and drought tolerance can result in negative covariation among leaf and root traits in improved *Silphium* seedlings (Gonzalez-Paleo et al., 2024). Understanding covariation among traits is key to domesticating perennial grains and oilseeds that yield sufficiently and maintain robust root systems. A remaining question in *Silphium* and other species is the extent to which water availability, and specifically the location of the primary source of water (surface, deeper in the soil), impacts resource allocation and trait covariation.

Research on trait covariation in plants often focuses on whole-organism response to availability or limitation of a specific resource. At least two aspects of water availability, water deficit and the location of available water, have consequences for aboveground biomass, root distribution, and root morphology (e.g., Fry et al., 2018; Gao et al., 2019; Zhou et al., 2018). It is well known that aboveground biomass is influenced by water deficit (Gao et al., 2019; Zollinger et al., 2006). Variation in aboveground responses of plants to water deficit may be attributable to differences in their root distributions, which also vary as a function of water deficit (Manschadi et al., 2006; Vadez et al., 2013) and the location of available water (Fry et al., 2018; Guevara et al., 2010). For example, when there is little water in the soil, roots may proliferate near the soil surface to capture intermittent precipitation (a shallow-rooted strategy) and/or grow deeper to reach water deeper in the soil (a deep-rooted strategy; Burridge et al., 2020; Manschadi et al., 2006; Schenk & Jackson, 2002). The root systems of many different species become either more shallow-rooted or more deep-rooted in response to water deficit (Manschadi et al., 2006); however, some species balance shallow-and deep-rooted distribution strategies with dimorphic root systems composed of both a mesh of fine roots near the soil surface and deep tap roots (Burridge et al., 2020; Ho et al., 2005). It is unclear how the balance of dimorphic root systems shift as the location of available water changes.

In addition to affecting the distribution of roots in the soil, water deficit and the location of available water impact root morphology (Kou et al., 2022; Zhou et al., 2018). Specifically, water deficit is associated with increased root diameter and decreased root length and root length density (Kou et al., 2022; Zhou et al., 2018). Water deficit is associated with a negative relationship between total root biomass and specific root length (i.e., the summed length of the roots divided by the summed mass of the roots; Zhou et al., 2018). When the location of available water is deeper in the soil, *Populus euphratica* seedlings tend to allocate more resources towards the roots than the shoots (Wang et al., 2015), and winter barley will produce a more dense root system near the soil surface (Ahmadi et al., 2018). Furthermore, roots closer to the soil surface can be more plastic in response to water location than roots deeper underground (Fry et al., 2018; Guevara & Giordano, 2015). While it is clear that water availability influences aboveground biomass, root distribution, and root morphology, how the location of available water (on the surface or deeper in the soil) influences covariation between aboveground and belowground plant structures is less well known.

In this study, we investigate the effects of the location of available water (on the surface or deeper in the soil) on aboveground and belowground traits and their relationships to one another. This study leverages a hybrid *Silphium* population derived from two ecologically and morphologically distinct *Silphium* species, *S. integrifolium* and *S. perfoliatum*, that differ in root distribution, morphology, and aboveground traits (Gonzalez-Paleo et al., 2023; Gonzalez-Paleo et al., 2024; Greve et al., 2023; Kowalski et al., 2004). We use this F1 backcross population in a greenhouse experiment to address how watering regime affects 1) aboveground and belowground biomass, 2) the distribution of the root system, 3) the relationship between aboveground mass and the distribution of the root system, and 4) the relationship between aboveground mass and the morphology of the root system. Expanded understanding of these relationships may be informative for the domestication and improvement of herbaceous, perennial crop candidate species that are high-yielding with root systems that provide key belowground ecosystem services.

## MATERIALS AND METHODS

### Study system

Two hundred seedlings (approximately one week in age) derived from an F1 backcross between the hybrid *S. integrifolium* x *S. perfoliatum* genotype 3ZZP and the *S. integrifolium* genotype BAD ASTRA were obtained from The Land Institute (Salina, KS). 3ZZP is the product of a cross between the *S. integrifolium* genotype BAD ASTRA (mother) and the *S. perfoliatum* genotype 1JHW (father). 1JHW was collected from a wild *S. perfoliatum* population in Antelope County, Nebraska, USA. BAD ASTRA was an *S. integrifolium* individual selected from the 2017 breeding population of *S. integrifolium* at The Land Institute, which had undergone several cycles of selection for feminization, seed size, and rust resistance. The original wild *S. integrifolium* population from which this breeding population originates is in Kansas, USA (David Van Tassel, pers. comm.). Seedlings from the F1 backcross population were sent to the Danforth Center in small plugs.

### Planting and greenhouse conditions

Upon arrival, plugs were transplanted to 8.94 cm wide pots (#700035C from T.O. Plastics; Otsego, Minnesota, USA) and kept outside, uncovered, for up to two weeks. Plants were then moved to a growth chamber for two weeks under the following conditions: 450 µmol of light, 50% humidity, 12:12 photoperiod; daytime temperature 30°C, nighttime temperature 21°C. After two weeks, plants were moved from the growth chamber to the greenhouse (50% humidity, 14:10 photoperiod; daytime temperature 28°C; nighttime temperature 22°C). After at least five weeks in the greenhouse, three cohorts of plants were transplanted at different timepoints into 22.7 L tree pots (approximately 30 cm wide by 60 cm deep) containing a mixture of 40% turface (Profile Products LLC, Buffalo Grove, Illinois, USA) / 30% sand (Quikrete, Atlanta, Georgia, USA) / 30% vermiculite (Therm-O-Rock East Inc., New Eagle, Pennsylvania, USA). Osmocote fertilizer (5 mL, 14:14:14 NPK; ICL Specialty Products, St. Louis, Missouri, USA) was placed in each large pot after every 12.7 cm of mixed media (approximately 20 mL of fertilizer in total).

### Experimental design

Plants (n = 144) were split randomly into three groups (cohorts) of 48 individuals per cohort. The three cohorts would spend different amounts of time growing in large tree pots (70 days, 50 days, or 30 days). This design allowed us to destructively harvest plants that had been growing in large tree pots for different amounts of time at approximately the same chronological age. The 70-day cohort was transplanted to the large tree pots 71 days after the plants arrived at the Danforth Center and spent approximately 70 days in the tree pots. The 50-day cohort was transplanted to the large tree pots on day 85 after arrival and spent approximately 50 days in the large tree pots. The 30-day cohort was transplanted to the large tree pots between days 104 and 108, spending approximately 30 days in the large tree pots. Plants that spent more time in the large tree pots also spent less time in the small pots before transplant. Destructive harvesting of all plants occurred between 131 and 141 days after arrival (Figure 1).

**Figure 1.**
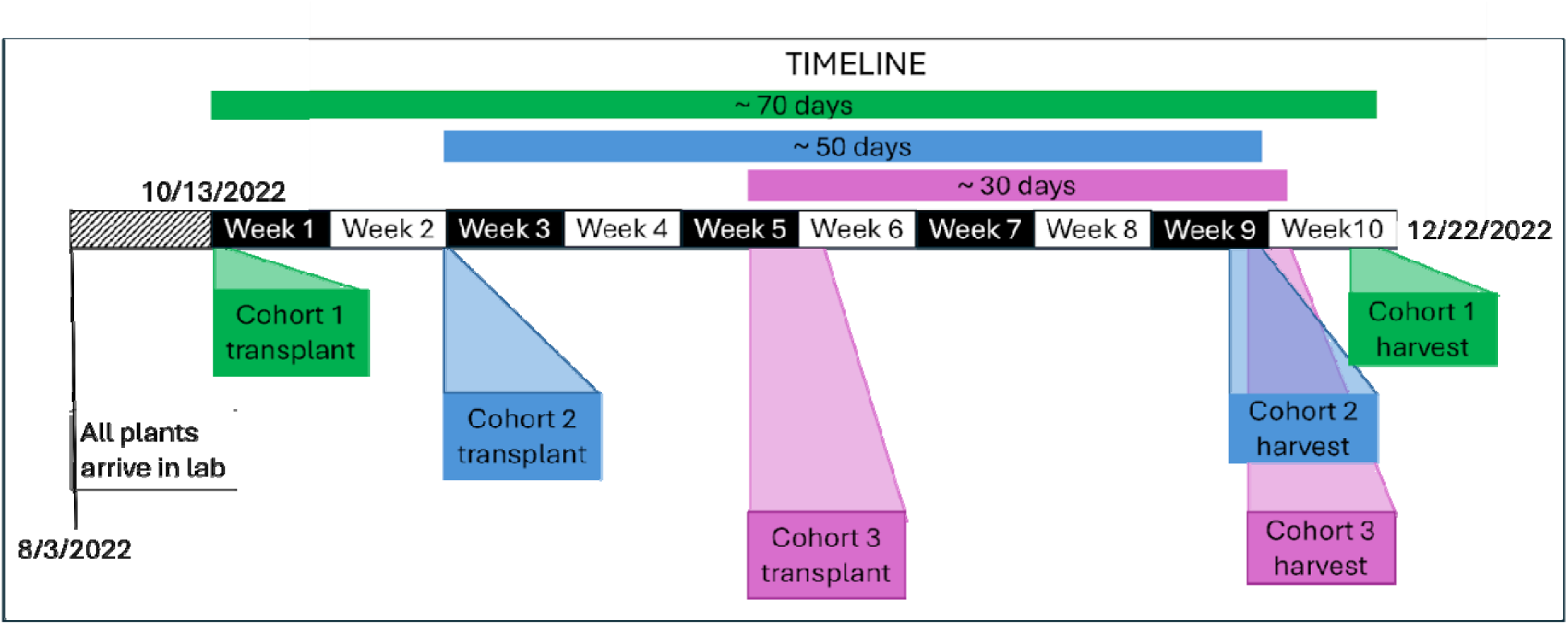
Timeline representing the timing of transplants and harvests for this experiment. The transplants were spaced in time so that the plants could be harvested at approximately the same chronological age after being in the large tree pots for different amounts of time. Each white or black segment of the timeline represents one week.

For the first seven days after transplant into large tree pots, all plants were watered from the top as needed with a hose. After seven days, the 48 plants within each cohort were split into two groups of 24 and assigned to receive either top-watering or bottom-watering treatments. At this point, the top-watered plants continued to be watered from the top, while the bottom-watered plants were placed, two pots per tray, in a 6.2 cm deep tray (#710245C from T.O. Plastics) and watered by filling the tray at least every two days until harvest. During harvest, we confirmed that pots with plants in different experimental groups contained approximately the same amount of water by comparing the weights of the pots. Pots were weighed individually in the greenhouse using a commercial scale (Rubbermaid; Winchester, Virginia, USA) with a maximum of 150 kg and a precision of +/-0.02 kg

#### Shoot phenotyping

Immediately prior to harvest, we recorded the developmental stage of the plants using guidelines provided by The Land Institute (Salina, Kansas, USA). This developmental stage scale ranges from 0.0 (seed) to 10.0 (completely senesced aboveground), where the beginning of bolting corresponds to a score of 2.1 and the beginning of flowering corresponds to a score of 5.1. Harvesting of aboveground biomass took place between 131 and 141 days after the plants arrived at the Danforth Center. All aboveground material was removed from the plant using pruning shears. Fresh aboveground biomass was measured in grams using a tabletop scale with a precision of +/-5 mg. To measure weight of dried aboveground biomass, aboveground biomass for each individual was placed in paper bags, dried at 37°C for at least six days, and then reweighed using the same scale.

#### Root phenotyping

Following harvest of aboveground biomass, we emptied each pot onto a greenhouse table covered with a fine mesh screen and washed the remaining substrate off the roots with a water hose. After removing most of the substrate and before any subsequent measurements, roots were divided into three portions: 1) pre-transplant root ball; 2) post-transplant upper roots; and 3) post-transplant lower roots (Figure 2). First, the plants retained a clearly observable “root ball” that formed pre-transplant (i.e., the portion of the root system that developed in the 8.94 cm wide pots prior to the transplant to the large tree pots). These pre-transplant root balls were easily distinguishable from roots that grew following transplant to the large tree pots: their root structure was unique, as was the surrounding substrate (Figure 2B). Second, immediately outside of the pre-transplant root balls, we observed post-transplant upper roots, roots that were within 20.32 cm of the top of the root system but outside of the pre-transplant root ball. This cutoff separates roots in the upper one-third of the pot from the roots in the lower two-thirds of the pot. Third, we observed roots that had grown the longest, which we refer to as the post-transplant lower roots. Post-transplant lower roots were those observed beyond 20.32 cm from the top of the root system. These roots were separated from the post-transplant upper roots by cutting them with scissors. Following removal, pre-transplant root balls, post-transplant upper roots, and post-transplant lower roots were placed in separate plastic bags and kept in a freezer at -20°C.

**Figure 2.**
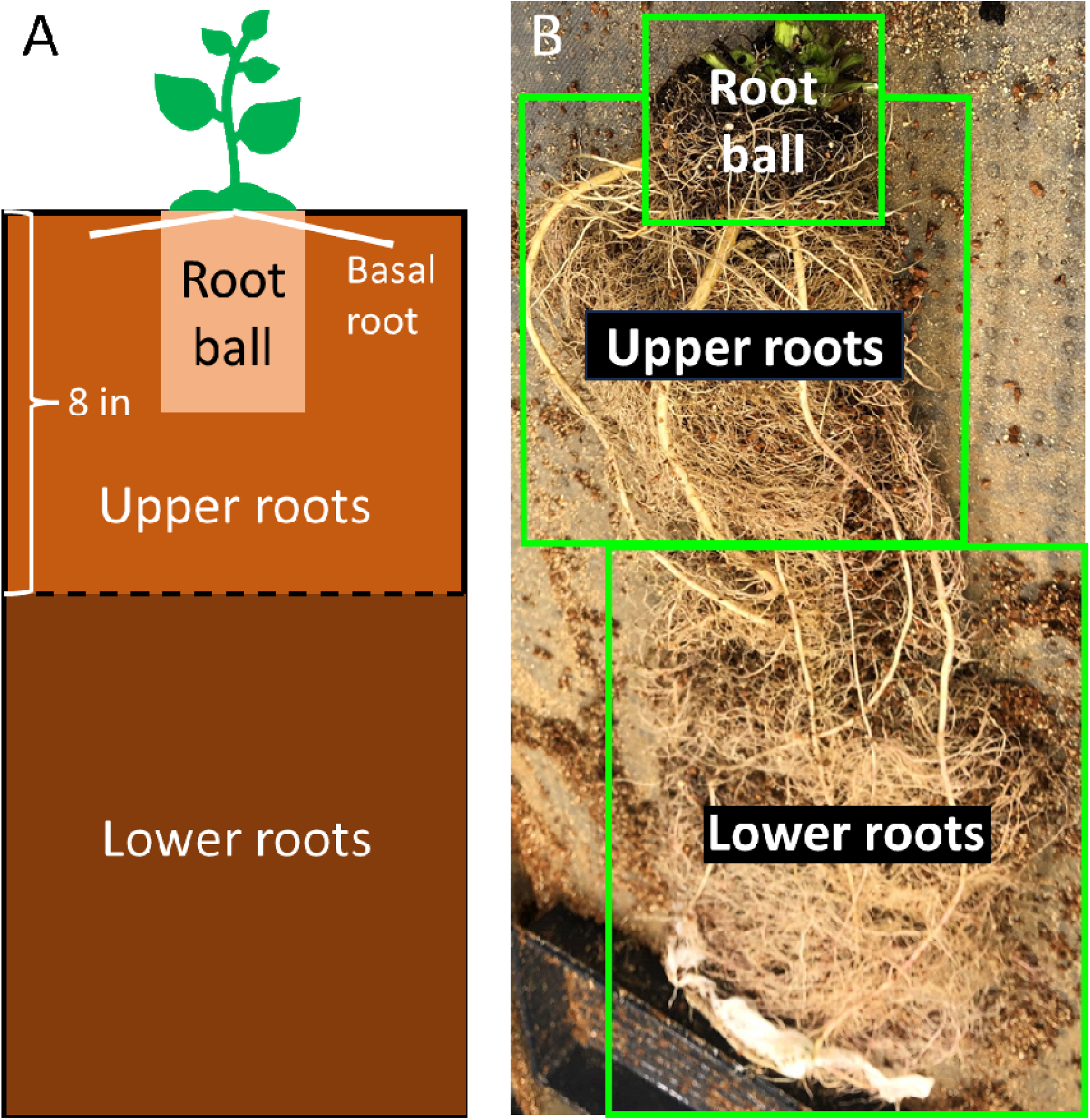
(A) Illustration of the location of basal roots and the three different root samples taken from each plant during harvest (i.e., pre-transplant root balls, post-transplant upper roots, and post-transplant lower roots). (B) Example root system during harvest with pre-transplant root balls, post-transplant upper roots, and post-transplant lower roots marked with green rectangles.

Pre-transplant root balls, post-transplant upper roots, and post-transplant lower roots were processed by washing off any remaining substrate, weighing cleaned roots, placing cleaned roots into a paper bag, drying cleaned roots at 60°C for at least three days, and then weighing the cleaned, dried roots. Total belowground mass was computed as the sum of pre-transplant root ball mass, post-transplant upper roots mass, and post-transplant lower roots mass. Root-shoot ratio was computed as total dried belowground mass divided by dried aboveground mass. In addition, for the pre-transplant root balls, we counted the number of basal roots (roots emerging from the base of the stem, just above where the root system begins) and measured the thickness of the three thickest basal roots with digital calipers before drying them. The basal roots emerged directly from the hypocotyl immediately above the root system (i.e., adventitiously) in the pre-transplant root ball samples and were relatively more coarse than the rest of the root system.

For the post-transplant lower roots, the three thickest post-transplant lower roots were trimmed to remove any lateral branches and imaged separately using an Epson 12000 XL scanner in color against a white background. A clear, plastic, water tray was placed on the scanner and filled approximately halfway with DI water. The root was then placed into the water and the lid was closed over the water tray. Images were scanned with a resolution of 600 dpi and saved locally as JPEG files. After scanning these individual roots, we weighed them separately.

Scanned images of individual roots from the post-transplant lower root samples were processed using RhizoVision Explorer 2.0.3 (Seethapalli et al., 2021). Using RhizoVision in batch mode, we removed root segments that were thinner than 1.25 mm. We increased the edge smoothing parameter from a default value of 2 to a value of 10 to smooth over the bases of any physically removed branches. The length of the remaining root as calculated by RhizoVision was then divided by the estimated dried mass of the root to obtain a measure of specific root length. Specific root length for each of the three individual roots were averaged to obtain mean specific root length.

### Analysis

All statistical analyses were conducted in RStudio (version 2023.12.0) with R (version 4.3.2; R Core Team, 2024). The packages *car* (version 3.1.3; Fox & Weisberg, 2019), *Hmisc* (version 5.2-3; Harrell Jr, 2025), *MASS* (version 7.3-60; Venables & Ripley, 2002), *olsrr* (version 0.6.1; Hebbali, 2024), and *sfsmisc* (version 1.1-20; Maechler, 2024) were used to conduct statistical analyses. The packages *ggplot2* (version 3.5.1; Wickham, 2016), *ggpubr* (version 0.6.0; Kassambara, 2023), *gridExtra* (version 2.3; Auguie, 2017), *gridGraphics* (version 0.5-1; Murrell & Wen, 2020), *patchwork* (version 1.3.0; Pederson, 2024), and *tidyr* (version 1.3.1; Wickham et al., 2024) were used to visualize results. One extreme outlier was removed from the bottom-watered condition for the 30-day cohort because of trait values that were beyond eight standard deviations from the mean trait value.

To address the question of how watering regime affects aboveground and belowground biomass, we compared top-and bottom-watered plants within each cohort using two-tailed Welch’s t-tests (Welch, 1947). Welch’s t-tests were selected because we observed unequal variances across watering conditions (Table S1). To test how watering regime influences the distribution of the root system (percent of root biomass of pre-transplant root balls and post-transplant lower roots), we compared top-and bottom-watered plants within each cohort using two-tailed Welch’s t-tests. To determine whether watering regime affects the relationship between aboveground mass and the distribution of roots, we conducted robust linear regressions separately for each cohort using the following models:

RootBallMass ∼ WateringRegime + AbovegroundMass + WateringRegime*AbovegroundMass

LowerRootsMass ∼ WateringRegime + AbovegroundMass + WateringRegime*AbovegroundMass

A significant interaction for either of these models would suggest that watering regime affects the relationship between the criterion variable (RootBallMass or LowerRootsMass) and aboveground mass for that cohort. Robust linear regressions were used instead of standard linear regressions to reduce the influence of outliers on the results (Table S2; Andersen, 2008). Finally, to address whether watering regime affects the relationship between aboveground mass and the morphology of the root system, we conducted robust linear regressions separately for each cohort using the following models:

NumberOfBasalRoots ∼ WateringRegime + AbovegroundMass + WateringRegime*AbovegroundMass

MeanBasalRootThickness ∼ WateringRegime + AbovegroundMass + WateringRegime*AbovegroundMass

MeanSpecificRootLength ∼ WateringRegime + AbovegroundMass + WateringRegime*AbovegroundMass

A significant interaction for any of these models would suggest that watering regime affects the relationship between the criterion variable (NumberOfBasalRoots, MeanBasalRootThickness, or MeanSpecificRootLength) and aboveground mass for that cohort.

## RESULTS

The hybrid *S. integrifolium* x *S. perfoliatum* F1 backcross population exhibited phenotypic variation in aboveground and belowground traits (Tables 1 and 2). Some of the plants had “cups” where the leaves met the stem, characteristic of *S. perfoliatum* (Gansberger et al., 2015). The plants also differed in leaf shape, phyllotaxy, and timing of bolting. For belowground traits, some plants had many thick basal roots and a small tap root, while others exhibited few basal roots and a large tap root. Watering regime affected both aboveground and belowground traits and their relationships to one another, as described below.

**Table 1.**
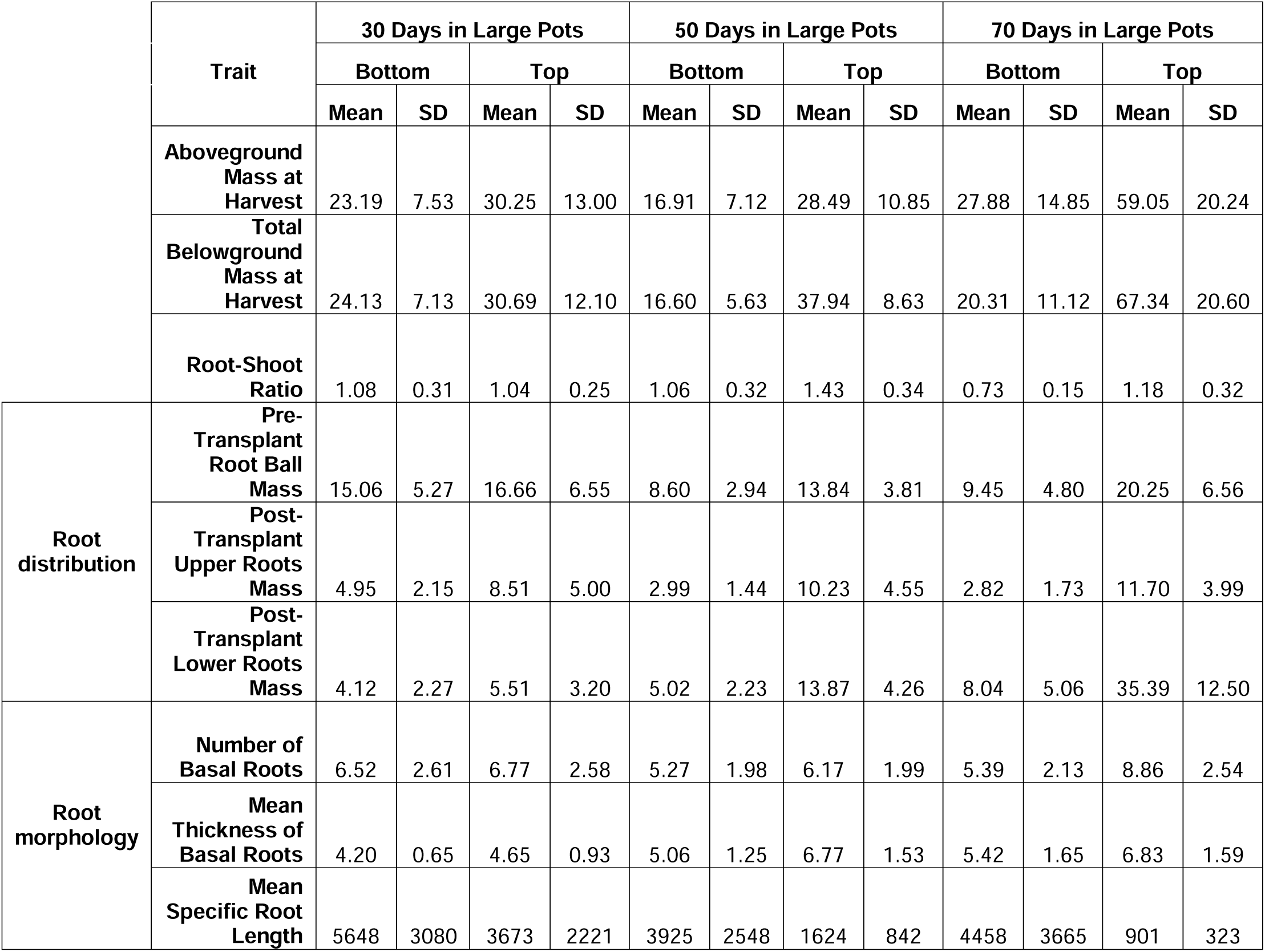
Descriptive statistics (means and standard deviations) for aboveground mass, total belowground mass, root-shoot ratio, pre-transplant root ball mass, post-transplant upper roots mass, post-transplant lower roots mass, number of basal roots, mean basal root thickness, and mean specific root length, separated by cohort and watering regime.

**Table 2.** Inferential statistics for Welch’s t-tests comparing top-and bottom-watered plants for each cohort in terms of aboveground mass, total belowground mass, root-shoot ratio, pre-transplant root ball mass, post-transplant upper roots mass, and post-transplant lower roots mass. Significant differences are highlighted with bold font. Aboveground and belowground traits vary significantly based on watering regime.

| Trait | Cohort | Welch's t | df | p |
| --- | --- | --- | --- | --- |
| Aboveground Mass | **30-day** | **2.22** | **33** | **0.034** |
|  | **50-day** | **3.19** | **18** | **0.005** |
|  | **70-day** | **5.78** | **36** | **< 0.001** |
| Total Belowground Mass | **30-day** | **2.20** | **34** | **0.035** |
|  | **50-day** | **7.40** | **18** | **< 0.001** |
|  | **70-day** | **9.30** | **30** | **< 0.001** |
| Root-Shoot Ratio | 30-day | 0.44 | 42 | 0.660 |
|  | **50-day** | **2.87** | **23** | **0.009** |
|  | **70-day** | **5.81** | **28** | **< 0.001** |
| Pre-Transplant Root Ball Mass | 30-day | 0.90 | 40 | 0.371 |
|  | **50-day** | **3.92** | **20** | **< 0.001** |
|  | **70-day** | **6.18** | **36** | **< 0.001** |
| Post-Transplant Upper Roots Mass | **30-day** | **3.08** | **28** | **0.005** |
|  | **50-day** | **5.30** | **13** | **< 0.001** |
|  | **70-day** | **9.41** | **27** | **< 0.001** |
| Post-Transplant Lower Roots Mass | 30-day | 1.67 | 38 | 0.102 |
|  | **50-day** | **6.51** | **16** | **< 0.001** |
|  | **70-day** | **9.35** | **26** | **< 0.001** |

### Watering regime affects aboveground and belowground biomass

Top-watered plants were significantly larger than bottom-watered plants as measured by aboveground mass, belowground mass, pre-transplant root ball mass, post-transplant upper roots mass, and post-transplant lower roots mass for 50-day and 70-day cohorts (Tables 1 and 2; Figures 3 and 4). Post-transplant upper roots mass also showed a difference between top-watered plants and bottom-watered plants for the 30-day cohort. Root-shoot ratio was significantly higher for top-watered plants than bottom-watered plants for 50-day and 70-day cohorts. Notably, the effect of watering regime on aboveground mass, pre-transplant root ball mass, post-transplant upper roots mass, and post-transplant lower roots mass increased with the amount of time the plants spent in the large tree pots.

**Figure 3.**
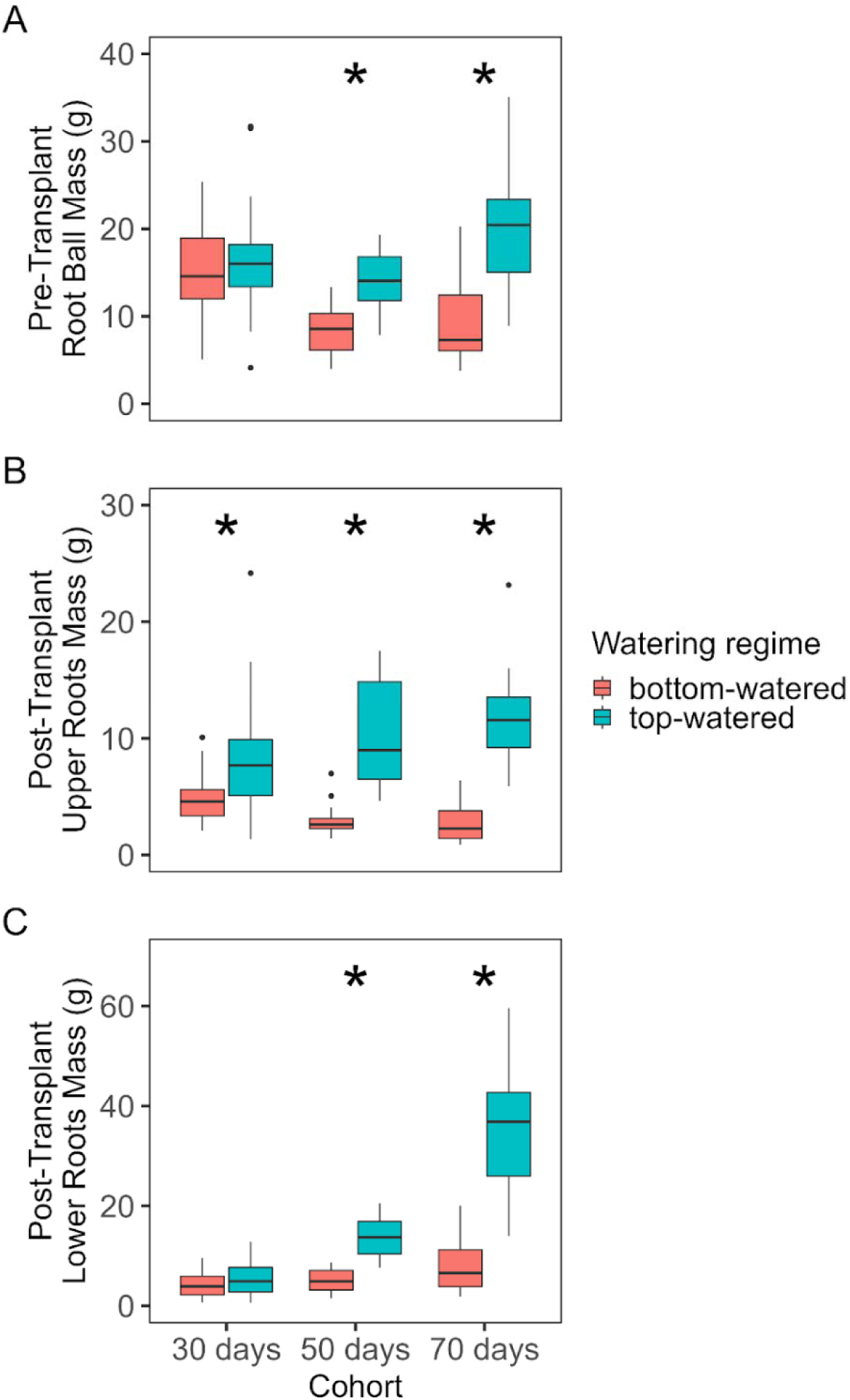
Box-plots representing the effects of watering regime for each cohort on (A) aboveground mass, (B) total belowground mass, and (C) root-shoot ratio. A black asterisk indicates a significant difference between top-and bottom-watered plants within that cohort. Aboveground biomass, belowground mass, and root-shoot ratio are significantly greater in top-watered plants.

**Figure 4.**
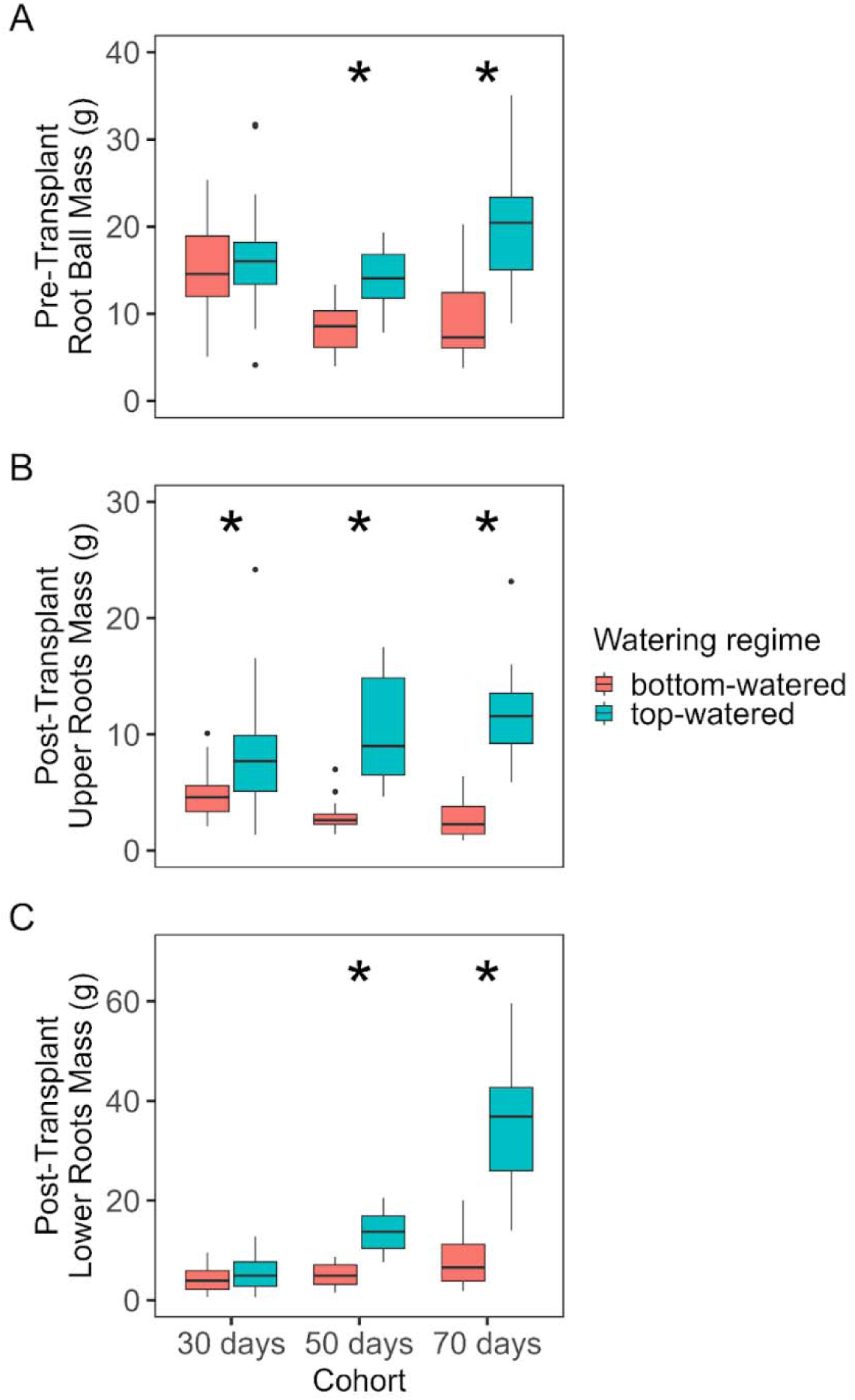
Box-plots representing the effects of watering regime for each cohort on (A) pre-transplant root ball mass, (B) post-transplant upper roots mass, and (C) post-transplant lower roots mass. A black asterisk indicates a significant difference between top-and bottom-watered plants within that cohort. Belowground biomass is greater in top-watered plants, but in some cases this effect is only observed in plants that have been in large tree posts for at least 50 days.

We considered the possibility that the watering regime affected the developmental stage of the plants and that differences in aboveground and belowground traits among water treatments reflected developmental differences. However, we did not find any evidence that watering regime affected developmental stage (all *p* > 0.05; Figure S1).

### Watering regime affects the distribution of roots

Top-watered plants distributed more root mass in the lower two-thirds of the pot than bottom-watered plants (Figure 5a, 5b). The proportion of the root system comprising the pre-transplant root ball was significantly higher for bottom-watered plants compared to top-watered plants for the 30-day (*t*(42) = 2.34, *p* = 0.024), 50-day (*t*(25) = 6.14, *p* < 0.001), and 70-day (*t*(34) = 10.50, *p* < 0.001) cohorts (Figure 5). The proportion of the root system comprising the post-transplant lower roots mass was significantly higher for top-watered plants compared to bottom-watered plants for the 50-day (*t*(19) = 2.20, *p* = 0.040) and 70-day (*t*(40) = 6.29, *p* < 0.001) cohorts. The effect of watering regime increased with time spent in the large tree pots.

**Figure 5.**
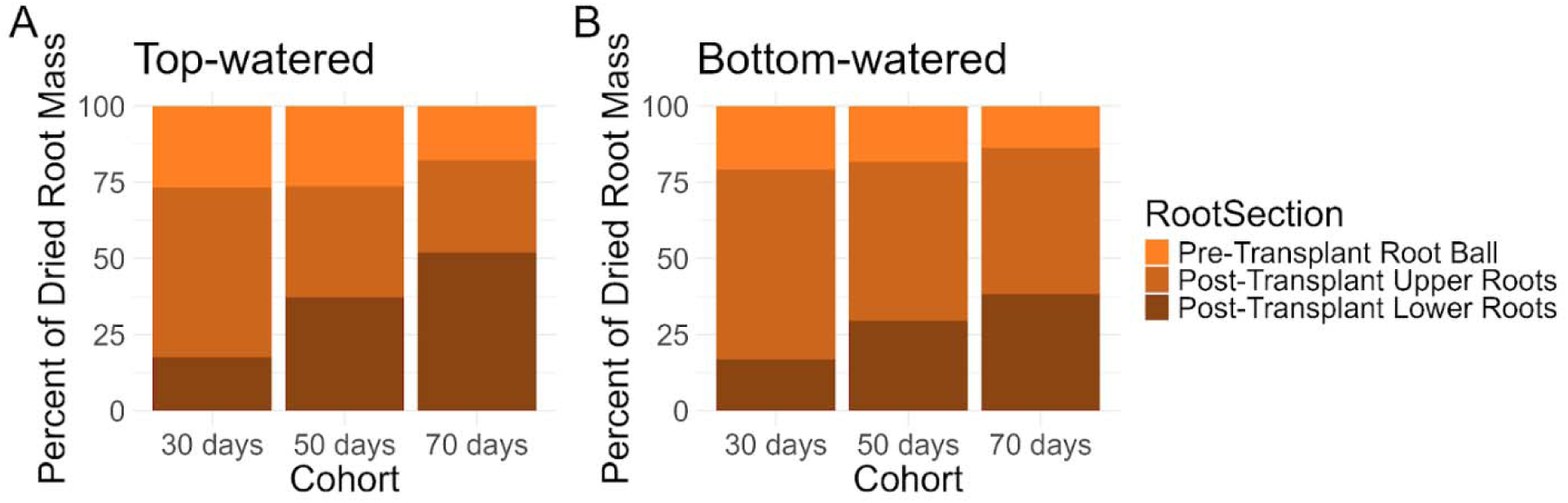
Stacked bar charts representing the distribution of dried root mass among pre-transplant root balls, post-transplant upper roots, and post-transplant lower roots for (A) top-watered and (B) bottom-watered plants within each cohort.

### Watering regime does not affect the relationship between aboveground mass and the distribution of the root system

Bigger plants were bigger everywhere: aboveground biomass was positively correlated with pre-transplant root ball mass and post-transplant lower roots. The relationship between aboveground biomass and pre-transplant root ball was similar for top-watered and bottom-watered plants. The relationship between aboveground biomass and post-transplant lower roots was also similar for top-watered and bottom-watered plants (Figure 6).

**Figure 6.**
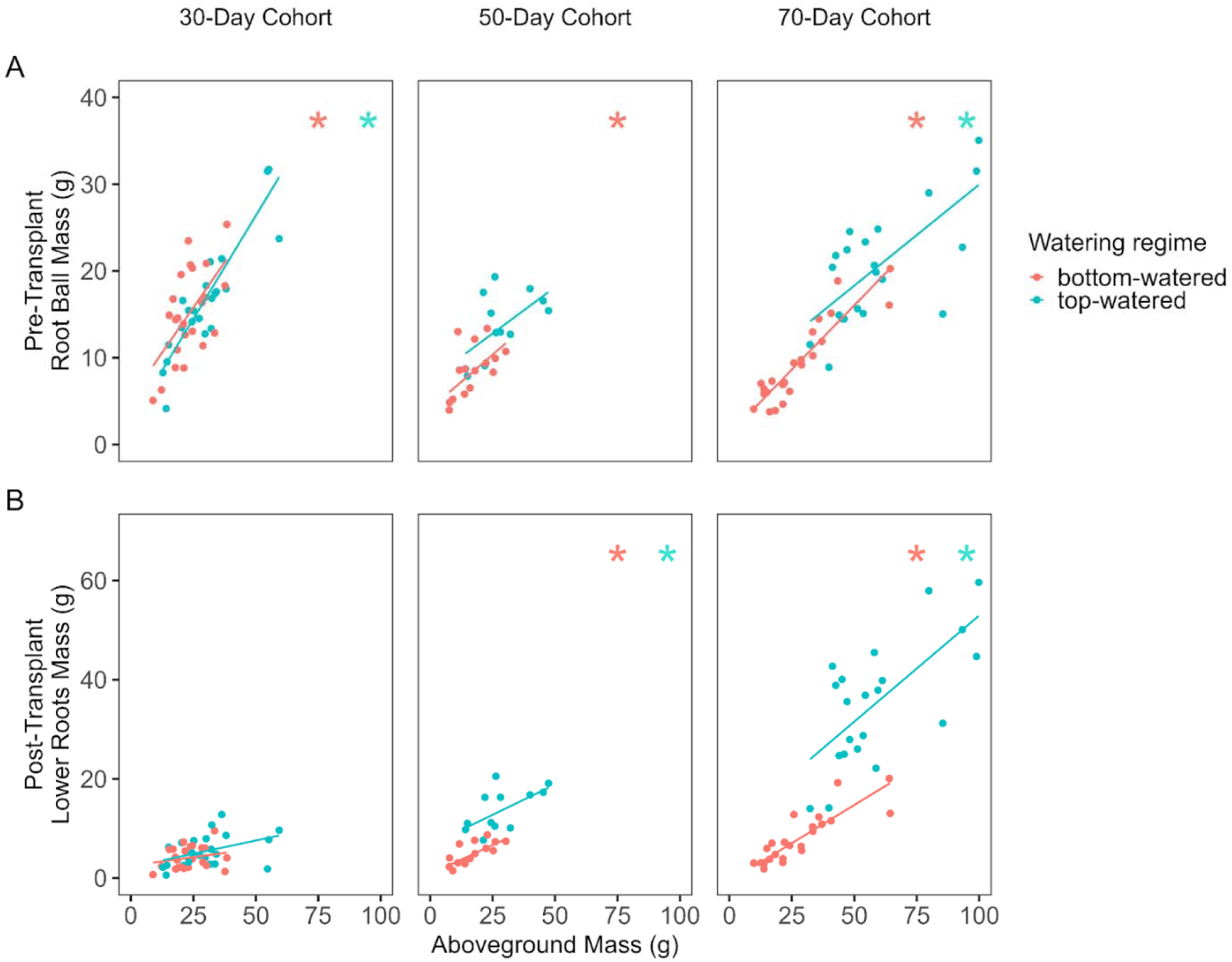
Scatterplots representing the relationships between (A) pre-transplant root ball mass and aboveground mass and (B) post-transplant lower roots mass and aboveground mass for each cohort. Lines of best fit within each scatterplot are derived from robust regressions. Pink asterisks indicate a significant correlation between the two variables for the bottom-watered plants within that cohort. Green asterisks indicate a significant correlation between the two variables for the top-watered plants within that cohort.

There were significant positive correlations between aboveground mass and pre-transplant root ball mass for top-watered plants (*r*(22) = 0.88, *p* < 0.001) and for bottom-watered plants (*r*(21) = 0.59, *p* = 0.003; Figure 6) for the 30-day cohort. For the 50-day cohort, there was a significant positive correlation between aboveground mass and pre-transplant root ball mass for the bottom-watered plants (*r*(13) = 0.56, *p* = 0.032). In addition, for the 50-day cohort, there were significant positive correlations between aboveground mass and post-transplant lower roots mass for top-watered plants (*r*(10) = 0.60, *p* = 0.038) and for bottom-watered plants (*r*(13) = 0.73, *p* = 0.002). For the 70-day cohort, there were significant positive correlations between aboveground mass and pre-transplant root ball mass for top-watered plants (*r*(19) = 0.68, *p* < 0.001) and for bottom-watered plants (*r*(21) = 0.91, *p* < 0.001), as well as significant positive correlations between aboveground mass and post-transplant lower roots mass for top-watered plants (*r*(19) = 0.68, *p* < 0.001) and for bottom-watered plants (*r*(21) = 0.85, *p* < 0.001).

### Watering regime affects the relationship between aboveground mass and the morphology of the root system

Although watering regime did not affect the relationship between aboveground biomass and the distribution of roots in the pot, we observed an interesting effect of watering regime on the relationship between aboveground biomass and root morphology. Aboveground biomass was negatively correlated with mean specific root length, and this relationship was strongest in the bottom-watered plants. Top-watered plants had lower mean specific root length than bottom-watered plants for 30-day (*t*(26) = 2.48, *p* = 0.018), 50-day (*t*(18) = 3.28, *p* = 0.004), and 70-day (*t*(22) = 4.63, *p* < 0.001) cohorts. For the 50-day cohort, there was a significant negative correlation between aboveground mass and mean specific root length for the bottom-watered plants (*r*(13) = -0.57, *p* = 0.026; Figure 7). For the 70-day cohort, there was a significant negative correlation between aboveground mass and mean specific root length for the top-watered plants (*r*(19) = -0.51, *p* = 0.019). For the 70-day cohort, there was also a significantly steeper negative relationship between aboveground mass and mean specific root length for the bottom-watered plants compared to the top-watered plants (*t*(40) = 2.06, *p* = 0.039).

**Figure 7.**
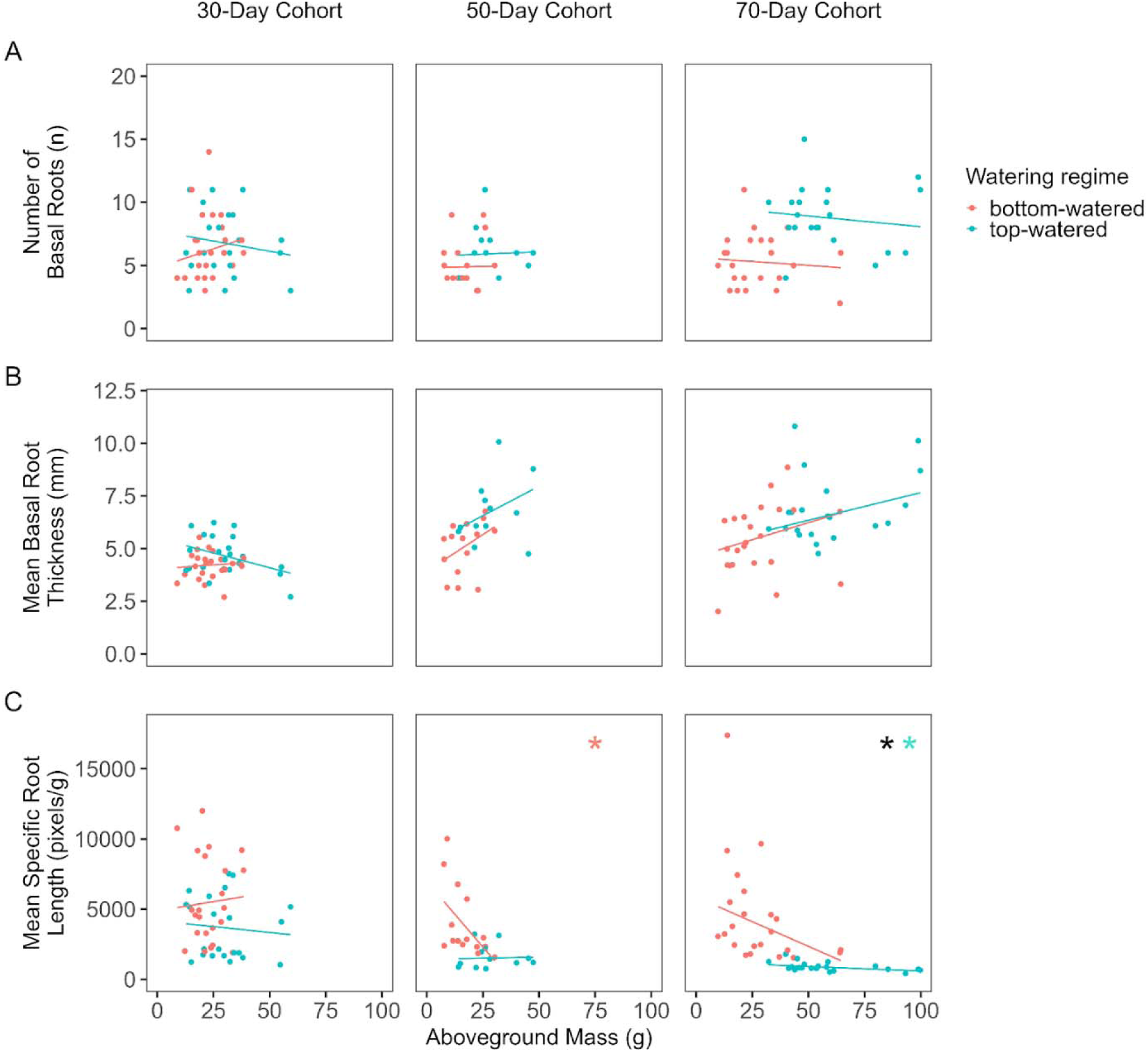
Scatterplots representing the relationships between (A) number of basal roots and aboveground mass, (B) mean basal root thickness and aboveground mass,and (C) mean specific root length and aboveground mass for each cohort. Lines of best fit within each scatterplot are derived from robust regressions. Pink asterisks indicate a significant correlation between the two variables for the bottom-watered plants within that cohort. Green asterisks indicate a significant correlation between the two variables for the top-watered plants within that cohort. Black asterisks indicate a significant difference between top-and bottom-watered plants for the relationship between that pair of variables for that cohort.

In addition, developmental stage moderated the effect of watering regime on mean specific root length for the 70-day cohort (*t*(40) = 4.70, *p* < .001). Specifically, the negative relationship between developmental stage and mean specific root length was steeper for bottom-watered plants compared to top-watered plants for the 70-day cohort (Figure S2).

## DISCUSSION

Through destructively harvested shoot and root systems that spent different amounts of time growing in large tree pots, this study provides insights into how plants allocate resources aboveground and belowground. Results show that the location of available water matters for overall plant growth both aboveground and belowground, consistent with previous studies (Fry et al., 2018; Guevara & Giordano, 2015; Wang et al., 2015). Compared to top-watered plants, bottom-watered plants were smaller aboveground and belowground (Figures 3 and 4). Bottom-watered plants distributed more root mass in the upper third of the pot and had higher mean specific root length (i.e., more length per mass) relative to top-watered plants. Furthermore, we observed that aboveground biomass was negatively correlated with mean specific root length and that this relationship was more negative in the bottom-watered plants. Below, we discuss how these results advance current understanding of whole-plant growth under different watering regimes and their potential implications for developing perennial herbaceous crops.

We expected that bottom-watered plants would produce more deep roots than the top-watered plants; however, we found that bottom-watered plants actually produced more root biomass in the upper portion of the pots than top-watered plants (Figure 5). At least two issues may have influenced the timing and extent of access to water in the bottom-watered plants. First, all pots were watered from the top prior to the experimental treatment. Once the watering treatment began, bottom-watered plants presumably had to grow roots further down into the pot to reach water, which likely required some amount of time. In contrast, the top-watered plants were able to use their already well-developed root system near the surface to access water starting at the beginning of the treatment. The relative delay in the timing of when roots are able to access water between bottom-watered plants and top-watered plants impacts not only how soon water can be accessed but also how much water can be absorbed. Second, beyond the timing of access to water, there was the difference in the extent of root mass available to actually access the water coming from the two different sources. In the bottom-watered plants, a limited proportion of the total root system grew towards water available in the tray; in contrast, a greater proportion of the total root mass was already concentrated towards the top of the pot, ready to access water delivered at the soil surface. For future work, researchers could estimate differences in the timing and extent of access to water delivered in different ways by estimating how much water the plant uptakes and by including another destructive sampling time point immediately before the bottom-watered plants are switched to having water available only at the bottom of the pots.

This finding challenges our intuitions about rooting strategies: top-watered plants grew relatively more roots into the bottom portion of the pot, and bottom-watered plants grew more roots than expected in the top portion of their pots. This pattern may be explained by previous observations that *Silphium* root systems are dimorphic. Specifically, we have seen large diameter, nearly vertical, deep, relatively permanent tap roots and fine, seasonal, nearly horizontal surface roots. Moisture limitation may stimulate the temporary production of surface roots, competing with resources needed for tap root production and leading to proportionately reduced root distribution into the lower pot. This study captured growth in pots over a few months; future work aimed at quantifying root growth over many years/months, outside of pots in natural conditions, will shed more light on relative patterns of growth among root types in species that exhibit dimorphic root morphology.

Another important observation of this study was that bottom-watered plants had a strong negative relationship between aboveground biomass and mean specific root length. Specifically, bottom-watered plants that were smaller aboveground grew deep roots with high mean specific root length (more length per unit of mass) than bottom-watered plants that were larger aboveground (Figure 7) and belowground. Previous studies have shown that under water deficit, specific root length is negatively correlated with belowground mass (Zhou et al., 2018). Across many different woody and herbaceous study systems, plants subject to water deficit tend to have higher specific root lengths and smaller belowground mass compared to control plants (Zhou et al., 2018). This pattern is consistent with the present study because, although we did not manipulate water deficit deliberately, bottom-watered plants appear to have had less access to water than top-watered plants.

While some of the bottom-watered plants appeared to prioritize building aboveground biomass, other bottom-watered plants seem to have prioritized exploration of the soil with higher mean specific root lengths. Differences among plants in response to the location of available water is likely a reflection of segregating variation in this population. Individuals from this population were not genetically identical and varied in the amount of introgression from *S. perfoliatum* in the *S. integrifolium* background. It is interesting to note that, even given this genetic variation, we found consistent effects of watering regime on aboveground biomass, root distribution, root morphology, and covariation among these traits.

There are at least two possible explanations for this combination of findings. First, the “economic” explanation is that *Silphium* balances the allocation of resources towards the acquisition of carbon (growing more aboveground biomass) with the acquisition of water (growing more roots; Reich, 2014). In the present study, the depth of available water changed after a week, and the bottom-watered plants appear to have responded by growing more explorative roots (i.e., higher mean specific root length) in the lower portions of the pots. This pattern was more pronounced for bottom-watered plants that were smaller aboveground compared to bottom-watered plants that were larger aboveground, possibly because smaller bottom-watered plants had more difficulty accessing water. According to previous research, shallow fibrous root systems tend to be more plastic than deep tap root systems in response to water deficit (Fry et al., 2018), and lateral roots near the soil surface tend to exhibit a more pronounced response to the location of water than roots at the end of the tap root (Guevara & Giordano, 2015). Root systems that can balance the production of shallow fibrous roots with the production of deep roots are considered dimorphic (Burridge et al., 2020; Ho et al., 2005), but it is unknown how dimorphic root systems respond to changes in the depth of available water. Future work can address the possibility that upper roots are more fibrous and absorptive than lower roots in dimorphic root systems such as those in *Silphium* by manipulating the location of available water and measuring water uptake and hydraulic conductance along the length of the root.

A second, “developmental” interpretation for the effects of watering regime on aboveground and belowground trait covariation is that top-watered plants were advanced developmentally compared to bottom-watered plants, despite being the same chronological age during harvest. Indeed, water deficit is known to delay phenology in annual crops such as maize (Sah et al., 2020) and wheat (Ihsan et al., 2016), as well as woody perennials (Adams et al., 2015). However, the relationship between developmental stage based on the timing of sexual reproduction and developmental changes in the secondary thickening of roots is unknown. Generally, we did not find evidence that watering regime the timing of sexual reproduction (Figure S1). However, we observed that sexual developmental stage moderated the effect of watering regime on mean specific root length for the 70-day cohort. Specifically, the negative relationship between sexual developmental stage and mean specific root length was steeper for bottom-watered compared to top-watered plants for the 70-day cohort. This finding is consistent with previous findings that specific root length tends to decrease with plant age in both perennial forbs (Pastor-Pastor et al., 2021) and trees (Li et al., 2021; Wang et al., 2024; Zheng et al., 2024). Although plant age and sexual developmental stage are most likely scaled differently, we would expect them to be correlated with each other and to be related to specific root length in similar ways.

One obvious limitation of this study is that the plants were grown in pots, which limited plant growth compared to what might have been observed in field conditions. According to Poorter et al. (2012), researchers should avoid ratios above 2 g/L and preferably have ratios under 1 g/L to minimize the effects of pot size on the availability of water, nutrients, and irradiance for photosynthesis. The pots used in this study were 22.7 L, which is much larger than pot sizes from most of the literature (Poorter et al., 2012), and had an average ratio of dry biomass to pot volume of 1.44 g/L. Future work should investigate whether pot size can affect covariation among aboveground biomass, root distribution, and root morphology in a hybrid *Silphium* population by comparing these results to those gathered from field-grown plants.

The effects of watering regime on covariation among aboveground mass, root distribution, and root morphology can inform the domestication of perennial sunflower relatives because future crops will need to balance yield with ecosystem services provided by their roots in response to varying environmental conditions and management practices such as irrigation. The ecosystem services provided by roots include carbon sequestration, the reduction of soil erosion, and the regeneration of soil (Crews et al., 2018). Considering root distribution and morphology during selection may help produce crops that sequester more carbon in the soil and prevent soil erosion (Wong et al., 2024). While a coarser root system (with lower specific root length) can allow for greater rooting depth and prevent soil erosion, a higher proportion of fine roots (with higher specific root length) can result in faster root turnover and better carbon sequestration (Wong et al., 2024). In addition, previous research has suggested that maximizing aboveground yield may not directly conflict with the carbon sequestration provided by roots because of positive genetic correlations between yield-related traits and the turnover of absorptive roots in switchgrass (Chen et al., 2021). Results described here provide evidence that watering regime affects the relationship between aboveground mass and specific root length. Future research should investigate whether the effects of watering regime on relationships among aboveground mass, root distribution, and root morphology have consequences for the ecosystem services provided by the roots of perennial crop candidates in potentially stressful environmental conditions such as low water availability.

Many crops have been bred to germinate and grow rapidly after sowing in order to out-compete weeds. Farmers often prepare the seed bed and plant in anticipation of sufficient soil moisture at the soil surface that is adequate for germination. However, after germination, the soil often quickly dries out, beginning at the surface. Therefore, successful seedlings must grow their roots downward faster than the advancing zone of soil desiccation. Our results here suggest that superficial irrigation or prolonged rainy conditions (simulated by the top-watered treatment) may be required for rapid, robust deep root growth in the majority of silphium seedlings. In contrast, conditions early in seedling development where the top of the soil profile is dry but adequate moisture remains below (simulated by the bottom-watered treatment) could trigger over-investment in shallow roots. While this root allocation strategy in response to soil drying may be adaptive in some natural environments, this strategy could lead to seedling mortality or slow stand establishment, outcomes that are unacceptable to farmers. Satisfactory stand establishment by direct seeding remains a challenge for *Silphium perfoliatum*, an otherwise attractive biomass crop in Europe (Shafer et al., 2015), and poor establishment of undomesticated species frequently limits prairie restoration (Applestein et al., 2018). We hypothesize that rapid deep root production regardless of moisture levels and gradients may be an important domestication trait for perennial crops.

As a possible next step, we suggest screening genetically diverse populations under water limiting or bottom-watered conditions for individuals with much greater than average investment in deep roots. It is encouraging to see that, under apparently water-limiting conditions (bottom-watered treatment), considerable phenotypic variation is evident among individuals after 70 days for lower roots mass (Figure 6B), basal root number (Figure 7A), and basal root thickness (Figure 7B). This was not a genetic study and the heritability of these traits cannot be estimated here. However, the fact that these seedlings showed a variety of combinations of aboveground traits from the parental species makes it reasonable to hypothesize that belowground traits in this population may also be segregating and recombining in new ways. These new combinations may allow artificial selection to remodel a root system that is adapted to neither of the native habitats of *S. integrifolium* or *S. perfoliatum* but, rather, to the novel environment created by farmers trying to cultivate perennial crops.

## CONCLUSIONS

We investigated how watering regime affects covariation among aboveground biomass, root distribution, and root morphology in a hybrid *Silphium* population. We found that bottom-watered plants were smaller aboveground. Their root systems were also smaller (lower total belowground mass), more superficial (greater proportion in the upper pot), and more exploratory (higher mean specific root length). Negative covariation between aboveground biomass and mean specific root length was also steeper for bottom-watered plants than top-watered plants. Presumably, bottom-watered plants had relatively more difficulty accessing available water, limiting their aboveground growth, because of the time required to grow towards the lower portions of the pots. To reach available water, bottom-watered plants grew more explorative roots (i.e., more length per mass). Differences among bottom-watered plants regarding whether they prioritized root exploration or the acquisition of biomass may be an indication of segregating variation in this population. Understanding the effects of the location of available water on aboveground and belowground trait covariation in a hybrid *Silphium* population can delimit which combinations of traits are possible in different environments and inform ongoing efforts to domesticate perennial members of the sunflower family.

## Acknowledgments

The authors thank Jack Braley, Zachary Harris, Danielle Hopkins, Maxwell Look, Shannon Meehan, Zoraya Piedra, Samantha Selby, Rachel Tavares, Isabella Vergara, Alex Windsor, Stella Woeltjen and other members of the Miller Lab for their help with the development of the research, sample processing, and providing comments on earlier drafts of the manuscript. The authors would like to thank the Donald Danforth Plant Science Center Plant Growth Facility (RRID:SCR_024902) staff for their help with plant care and maintenance of the greenhouse facilities. Funding for this research was provided by Saint Louis University, Donald Danforth Plant Science Center, Foundation for Food and Agriculture Research Seeding Solution -Next Generation Crops (CA20-SS-0000000123), and New Roots for Restoration Biology Integration Institute (NSF 2120153).

## Author contributions

Following a CRediT taxonomy (https://credit.niso.org), T.T. contributed to conceptualization, data curation, formal analysis, investigation, methodology, project administration, software, supervision, visualization, writing - original draft, and writing - review and editing. M.T.H. contributed to conceptualization, investigation, methodology, project administration, supervision, and writing - review and editing. M.M. contributed to investigation, methodology, and writing - review and editing. M.J.R. contributed to funding acquisition, methodology, and writing - review and editing. D.V.T. contributed to funding acquisition, resources, and writing - review and editing. A.J.M. contributed to conceptualization, funding acquisition, methodology, project administration, resources, supervision, and writing - review and editing.

## Data availability statement

All of the data and analysis scripts have been deposited to Zenodo (https://doi.org/10.5281/zenodo.17203240).

## Supplementary Information

**Table S1.** Test statistics and p-values for Levene’s test for the equality of variances, nonparametric Mann-Whitney-Wilcoxon tests, and Welch’s t-tests comparing bottom-watered and top-watered plants for each cohort. A significant p-value (*p* < 0.05) for Levene’s test indicates that bottom-watered and top-watered plants have unequal variances. In case of a significant Levene’s test, Mann-Whitney-Wilcoxon tests or Welch’s t-tests may be preferred over typical Student’s t-tests. For the present study, nonparametric tests do not change the overall pattern of results. Significant p-values are highlighted with bold font.

| Trait | Cohort | Levene |  | Mann-Whitney-Wilcoxon |  | Welch's t-test |  |
| --- | --- | --- | --- | --- | --- | --- | --- |
|  |  | F | p | W | p | t | p |
| Aboveground Mass | 30-day | 3.29 | 0.077 | 170.0 | 0.061 | 2.22 | **0.034** |
|  | 50-day | 1.77 | 0.196 | 33.0 | **0.004** | 3.19 | **0.005** |
|  | 70-day | 1.72 | 0.196 | 42.0 | **< 0.001** | 5.78 | **< 0.001** |
| Total Belowground Mass | 30-day | 3.31 | 0.076 | 174.0 | 0.075 | 2.20 | **0.035** |
|  | 50-day | 3.32 | 0.080 | 2.0 | **< 0.001** | 7.40 | **< 0.001** |
|  | 70-day | 7.13 | **0.011** | 9.0 | **< 0.001** | 9.30 | **< 0.001** |
| Root-Shoot Ratio | 30-day | 0.29 | 0.590 | 256.0 | 0.955 | 0.44 | 0.660 |
|  | 50-day | 0.00 | 0.964 | 40.0 | **0.014** | 2.87 | **0.009** |
|  | 70-day | 8.56 | **0.006** | 42.0 | **< 0.001** | 5.81 | **< 0.001** |
| Pre-Transplant Root Ball Mass | 30-day | 0.15 | 0.704 | 218.5 | 0.440 | 0.90 | 0.371 |
|  | 50-day | 1.49 | 0.234 | 28.0 | **0.002** | 3.92 | **< 0.001** |
|  | 70-day | 1.44 | 0.237 | 43.5 | **< 0.001** | 6.18 | **< 0.001** |
| Post-Transplant Upper Roots Mass | 30-day | 5.49 | **0.024** | 120.5 | **0.003** | 3.08 | **0.005** |
|  | 50-day | 21.48 | **< 0.001** | 5.0 | **< 0.001** | 5.30 | **< 0.001** |
|  | 70-day | 8.21 | **0.006** | 2.0 | **< 0.001** | 9.41 | **< 0.001** |
| Post-Transplant Lower Roots Mass | 30-day | 2.90 | 0.096 | 184.0 | 0.121 | 1.67 | 0.102 |
|  | 50-day | 16.21 | **< 0.001** | 1.0 | **< 0.001** | 6.51 | **< 0.001** |
|  | 70-day | 13.10 | **< 0.001** | 4.0 | **< 0.001** | 9.35 | **< 0.001** |

**Table S2.** Test statistics and p-values for Shapiro-Wilk and Kolmogorov-Smirnov tests for the normality of residuals for each regression model and each cohort. Significant p-values (*p* < 0.05) for both tests indicate that the residuals of a linear regression model deviate from normality. Out of an abundance of caution, robust regressions were used to reduce the influence of outliers on the results. Significant p-values are highlighted in bold font.

| Criterion variable | Predictors | Cohort | Shapiro-Wilk |  | Kolmogorov-Smirnov |  |
| --- | --- | --- | --- | --- | --- | --- |
|  |  |  | W | p | D | p |
| Pre-Transplant Root Ball Mass | Aboveground Mass x Watering Regime | 30-day | 0.978 | 0.558 | 0.070 | 0.969 |
|  | Aboveground Mass x Watering Regime | 50-day | 0.883 | **0.006** | 0.215 | 0.141 |
|  | Aboveground Mass x Watering Regime | 70-day | 0.967 | 0.236 | 0.095 | 0.787 |
| Post-Transplant Lower Roots Mass | Aboveground Mass x Watering Regime | 30-day | 0.970 | 0.296 | 0.111 | 0.595 |
|  | Aboveground Mass x Watering Regime | 50-day | 0.938 | 0.108 | 0.147 | 0.555 |
|  | Aboveground Mass x Watering Regime | 70-day | 0.968 | 0.263 | 0.112 | 0.598 |
| Number of Basal Roots | Aboveground Mass x Watering Regime | 30-day | 0.958 | 0.104 | 0.100 | 0.726 |
|  | Aboveground Mass x Watering Regime | 50-day | 0.910 | **0.022** | 0.172 | 0.358 |
|  | Aboveground Mass x Watering Regime | 70-day | 0.978 | 0.572 | 0.084 | 0.891 |
| Mean Basal Root Thickness | Aboveground Mass x Watering Regime | 30-day | 0.986 | 0.868 | 0.063 | 0.989 |
|  | Aboveground Mass x Watering Regime | 50-day | 0.970 | 0.594 | 0.108 | 0.880 |
|  | Aboveground Mass x Watering Regime | 70-day | 0.975 | 0.457 | 0.091 | 0.832 |
| Mean Specific Root Length | Aboveground Mass x Watering Regime | 30-day | 0.934 | **0.014** | 0.167 | 0.145 |
|  | Aboveground Mass x Watering Regime | 50-day | 0.962 | 0.403 | 0.122 | 0.776 |
|  | Aboveground Mass x Watering Regime | 70-day | 0.737 | **< 0.001** | 0.220 | **0.024** |

**Figure S1.**
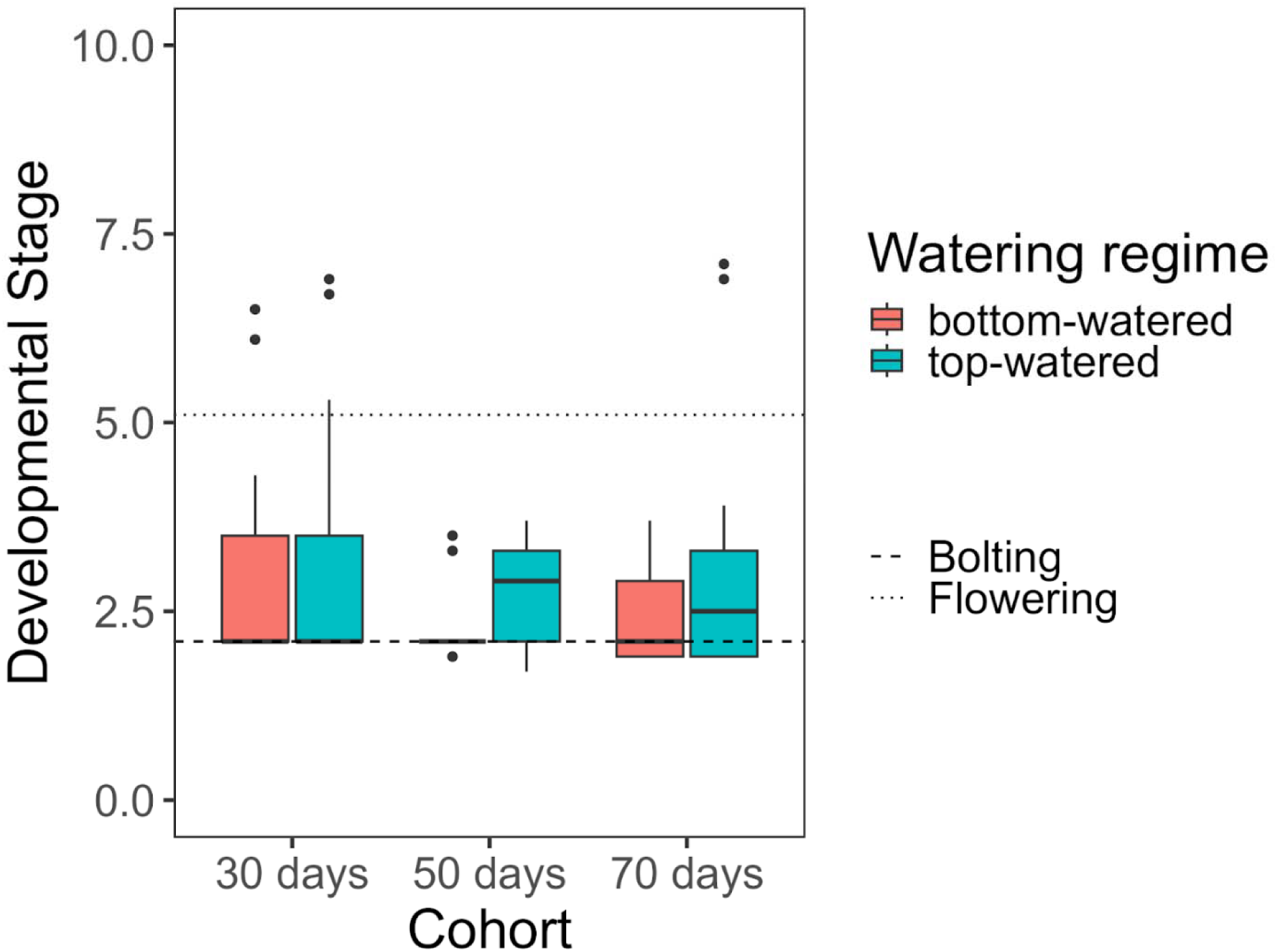
Box-plot representing the effects of watering regime for each cohort on developmental stage. The dashed line represents the developmental stage at which the plants began bolting, and the dotted line represents the developmental stage at which the plants began flowering. For each cohort, we compared developmental stages in top-watered and bottom-watered plants using a Welch’s t-test and did not find any significant differences (all *p*s > .05).

**Figure S2.**
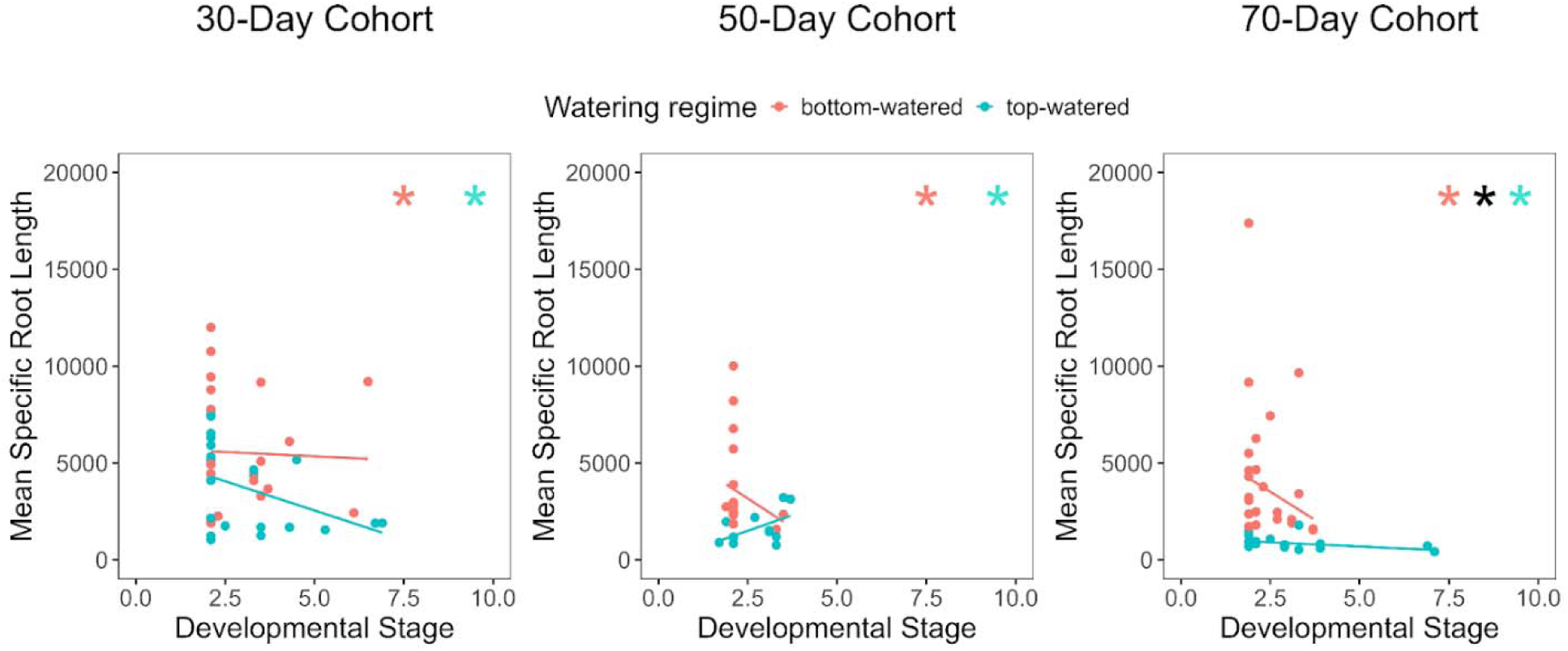
Scatterplots representing the relationships between developmental stage and mean specific root length for each cohort. Lines of best fit within each scatterplot are derived from robust regressions. Pink asterisks indicate a significant correlation between the two variables for the bottom-watered plants within that cohort. Green asterisks indicate a significant correlation between the two variables for the top-watered plants within that cohort. Black asterisks indicate a significant difference between top-and bottom-watered plants for the relationship between that pair of variables for that cohort.

